# Evidence of antigenic drift in the fusion machinery core of SARS-CoV-2 spike

**DOI:** 10.1101/2023.09.27.559757

**Authors:** Timothy J.C. Tan, Abby Odle, Ruipeng Lei, Kenneth A. Matreyek, Stanley Perlman, Lok-Yin Roy Wong, Nicholas C. Wu

## Abstract

Antigenic drift of SARS-CoV-2 is typically defined by mutations in the N-terminal domain and receptor binding domain of spike protein. In contrast, whether antigenic drift occurs in the S2 domain remains largely elusive. Here, we perform a deep mutational scanning experiment to identify S2 mutations that affect binding of SARS-CoV-2 spike to three S2 apex public antibodies. Our results indicate that spatially diverse mutations, including D950N and Q954H, which are observed in Delta and Omicron variants, respectively, weaken the binding of spike to these antibodies. Although S2 apex antibodies are known to be non-neutralizing, we show that they confer partial protection *in vivo*. We further demonstrate that such *in vivo* protection activity is diminished by the natural mutation D950N. Overall, this study indicates that the S2 domain of SARS-CoV-2 spike can undergo antigenic drift, which represents a potential challenge for the development of more universal coronavirus vaccines.

## INTRODUCTION

As the major antigen of severe acute respiratory syndrome coronavirus 2 (SARS-CoV-2), the spike (S) glycoprotein has undergone extensive antigenic drift since the beginning of the COVID-19 pandemic^1^. SARS-CoV-2 S protein is a homotrimer with an N-terminal domain (NTD), a receptor-binding domain (RBD), and an S2 domain. S protein facilitates virus entry by engaging the host receptor angiotensin-converting enzyme II (ACE2) via RBD and mediating virus-host membrane fusion through the fusion machinery in S2^2^. While all three domains in S can elicit antibody responses during infection or vaccination, the neutralizing potency of antibodies to RBD and NTD are typically much higher than those to S2^3^. Consistently, mutations in RBD and NTD are key determinants of SARS-CoV-2 antigenic drift^1,4,5^. Although mutations in S2 have also emerged in circulating SARS-CoV-2 variants^1^, they are thought to mainly affect the stability and fusogenicity of S protein^6–8^. As a result, whether S2 mutations play a role in the antigenic drift of SARS-CoV-2 remain largely elusive.

Due to the relatively high sequence conservation of S2, human antibodies to S2 can achieve exceptional breadth. For example, human antibodies to the S2 fusion peptide can neutralize coronavirus strains from different genera (α, β, γ and δ)^9–12^. Besides, human antibodies to the S2 stem helix can neutralize diverse β-coronavirus strains^13–17^. Additionally, a public clonotype to the apex of S2 can cross-react with multiple sarbecoviruses^18,19^. This public clonotype is encoded by IGHV1-69/IGKV3-11 with complementarity determining region (CDR) H3 and L3 lengths of 15 and 11 amino acids (IMGT numbering), respectively^18,19^. Although S2 antibodies usually have weak neutralizing activity, antibodies to fusion peptide and stem helix have been shown to confer *in vivo* protection against SARS-CoV-2 infection^9–17^. Given that S2 antibodies are commonly observed in both vaccinated and infected individuals^20,21^, they likely exert selection pressure on the circulating SARS-CoV-2.

In this study, we showed that the IGHV1-69/IGKV3-11 public clonotype to the apex of S2 confers partial *in vivo* protection, despite their lack of neutralizing activity^18^. Subsequently, a deep mutational scanning experiment was performed to probe the effects of S2 mutations on the cell-surface binding activity of three IGHV1-69/IGKV3-11 S2 antibodies, namely COVA1-07, COVA2-14, and COVA2-18. Specifically, we focused on single amino acid mutations within the first heptad repeat (HR1) and central helix (CH). Our results revealed that D950N and Q954H, which are observed in Delta and Omicron variants, respectively^1^, weakened binding of SARS-CoV-2 S to all three IGHV1-69/IGKV3-11 S2 antibodies. Additional *in vivo* experiment further suggested that protection mediated by COVA2-18 was weakened by D950N. Collectively, these results indicate that S2 mutations contribute to SARS-CoV-2 antigenic drift.

## RESULTS

### *In vivo* protection activity of IGHV1-69/IGKV3-11 S2 antibodies

Previous studies have reported a public clonotype against the S2 domain that is encoded by IGHV1-69/IGKV3-11^18,19^. Here, we tested the *in vivo* protection activity of three previously identified IGHV1-69/IGKV3-11 S2 antibodies, namely COVA1-07, COVA2-14, and COVA2-18^18^. Based on survival analysis (**Figure 1A**) and weight loss profiles (**Figure 1B**), all three antibodies showed prophylactic protection *in vivo* against a lethal challenge of SARS2-N501Y_MA30_, which is a mouse-adapted strain of SARS-CoV-2^22^. BALB/c mice treated with COVA1-07 showed 60% survival, while those with COVA2-14 and COVA2-18 showed 80% survival. In contrast, all mice treated with CR9114, which is an influenza virus-specific antibody^23^, succumbed to infection. This result indicates that although IGHV1-69/IGKV3-11 S2 antibodies have no neutralizing activity^18^, they contribute to protective antibody responses against severe SARS-CoV-2 infection. Similarly, a previous study has reported that COV2-2164, which is another member of the IGHV1-69/IGKV3-11 public clonotype to S2, modestly reduces the viral load in the lung and brain of K18-hACE2 transgenic mice upon SARS-CoV-2 infection, albeit with no effect on weight loss^19^.

**Figure 1.**
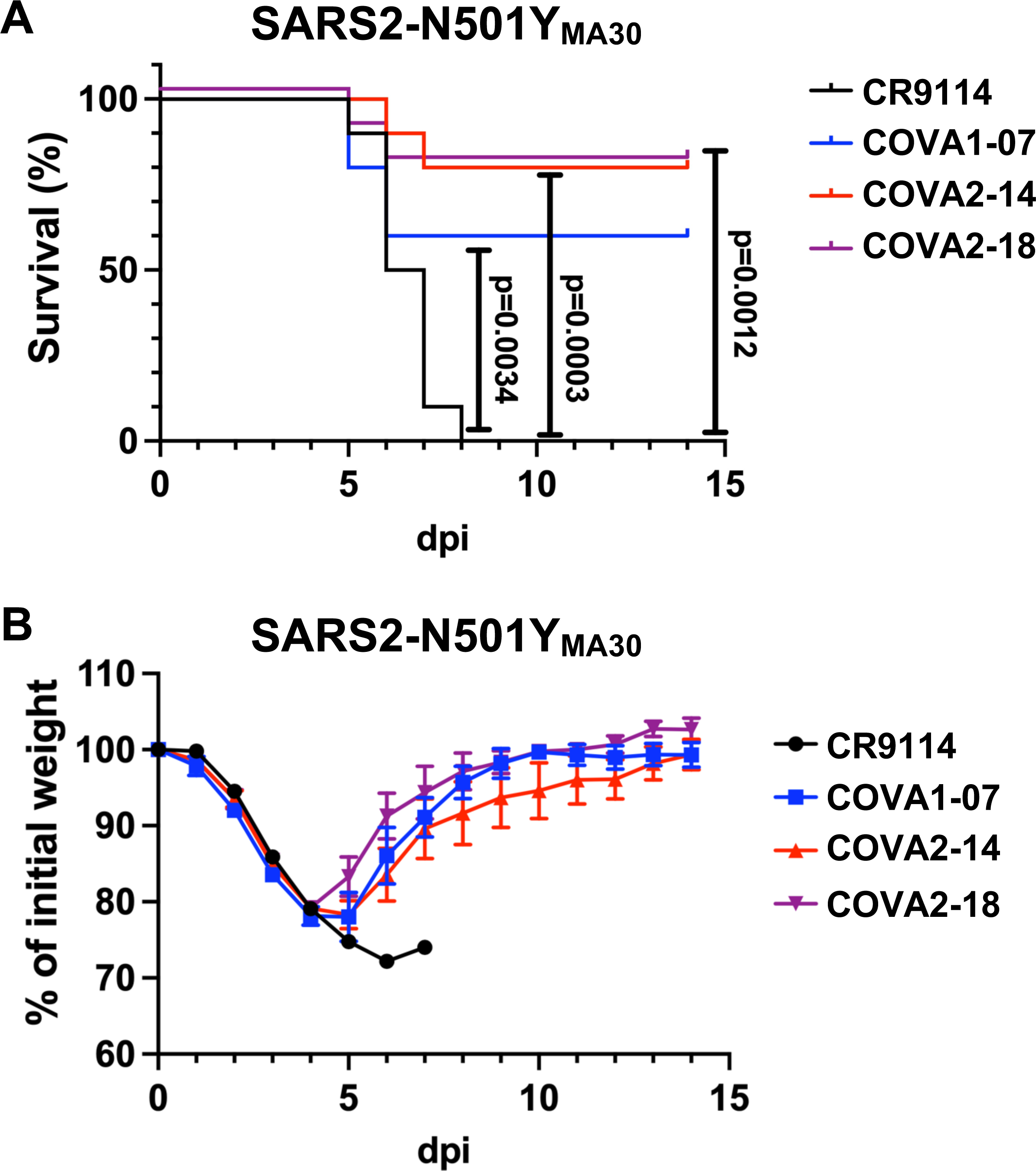
*In vivo* protection activity of IGHV1-69/IGKV3-11 S2 antibodies. BALB/c mice were intraperitoneally injected with 200 μg of CR9114 (black), COVA1-07 (blue), COVA2-14 (red), or COVA2-18 (purple) 16 hours before intranasal infection with 5,000 PFU of SARS2-N501Y_MA30_, which is a non-recombinant mouse-adapted strain of SARS-CoV-2^22^. Of note, non-recombinant SARS2-N501Y_MA30_ was used in this experiment. **(A)** Kaplan-Meier survival curves, and **(B)** weight loss profiles are shown (*n* = 10). Weight loss profiles are shown as mean ± SEM (standard error of the mean).

### Deep mutational scanning of S2 against IGHV1-69/IGKV3-11 antibodies

Given that IGHV1-69/IGKV3-11 S2 antibodies exhibited *in vivo* protection activity, we were interested in whether mutations in S2 could influence their binding activity. To systematically identify mutations on the S protein that affect binding to IGHV1-69/IGKV3-11 S2 antibodies, we performed a deep mutational scanning (DMS) experiment using our previously constructed S mutant library, which contained all possible amino acid mutations from residues 883 to 1034, spanning HR1 and CH in the S2 domain^24^. Briefly, this mutant library was displayed on human embryonic kidney 293T (HEK293T) landing pad cells. Fluorescence-activated cell sorting (FACS) was then performed to sort individual mutants according to their cell-surface binding activity to COVA1-07, COVA2-14, and COVA2-18 (**Figure S1**). Occurrence frequency of each mutant in different sorted populations was quantified by next-generation sequencing. For each experiment, a binding score was computed for each of the 1931 missense mutations, 122 silent mutations, and 132 nonsense mutations. The binding score was normalized such that the average score of silent mutations was 1 and that of nonsense mutations was 0.

Our DMS experiments were highly reproducible, with a Pearson correlation coefficient of 0.90 to 0.92 between independent biological replicates (**Figure S2A-C**). In addition, the binding score distributions of nonsense and silent mutations had minimal overlap, indicating that our DMS experiments could distinguish mutants with different cell-surface binding activities to COVA1-07, COVA2-14, and COVA2-18 (**Figure S2D-F**). We also observed a high Pearson correlation coefficient of 0.97 among the DMS experiments against COVA1-07, COVA2-14, and COVA2-18 (**Figure S2G-I**), indicating that the same mutation would have similar effect on cell-surface binding activity against these three antibodies. This result was not unexpected given that COVA1-07, COVA2-14, and COVA2-18 belong to the same IGHV1-69/IGKV3-11 public clonotype^18^.

### Two natural S2 mutations weaken binding to IGHV1-69/IGKV3-11 antibodies

To investigate if the binding activity of IGHV1-69/IGKV3-11 S2 antibodies could be affected by S2 mutations, we aimed to identify S2 mutations that weakened the binding activity to COVA1-07, COVA2-14, and COVA2-18. Since our deep mutational scanning experiments relied on cell surface display, mutations that lowered the cell-surface expression level would result in a lower binding score even if it did not affect the antibody binding affinity. Therefore, we compared the binding scores of individual mutations to their cell-surface expression levels which were previously quantified using the RBD antibody CC12.3^24,25^. As expected, there was a mild correlation between binding score and expression score, which is a proxy for cell-surface expression level^24^ (Pearson correlation coefficient of 0.48 to 0.49, **Figure 2**). Nevertheless, there were mutations with high expression scores but low binding scores, such as T961F, V987C, Q1005R, and Q1010W, all of which rarely occurred in circulating SARS-CoV-2^26^. Similarly, the low binding scores of D950N and Q954H, which are fixed in Delta and Omicron variants, respectively^1^, did not seem to be explained by the expression score alone. As a result, the low binding scores of these mutations were likely due to their effects on the antibody binding affinity, rather than cell-surface expression level.

**Figure 2.**
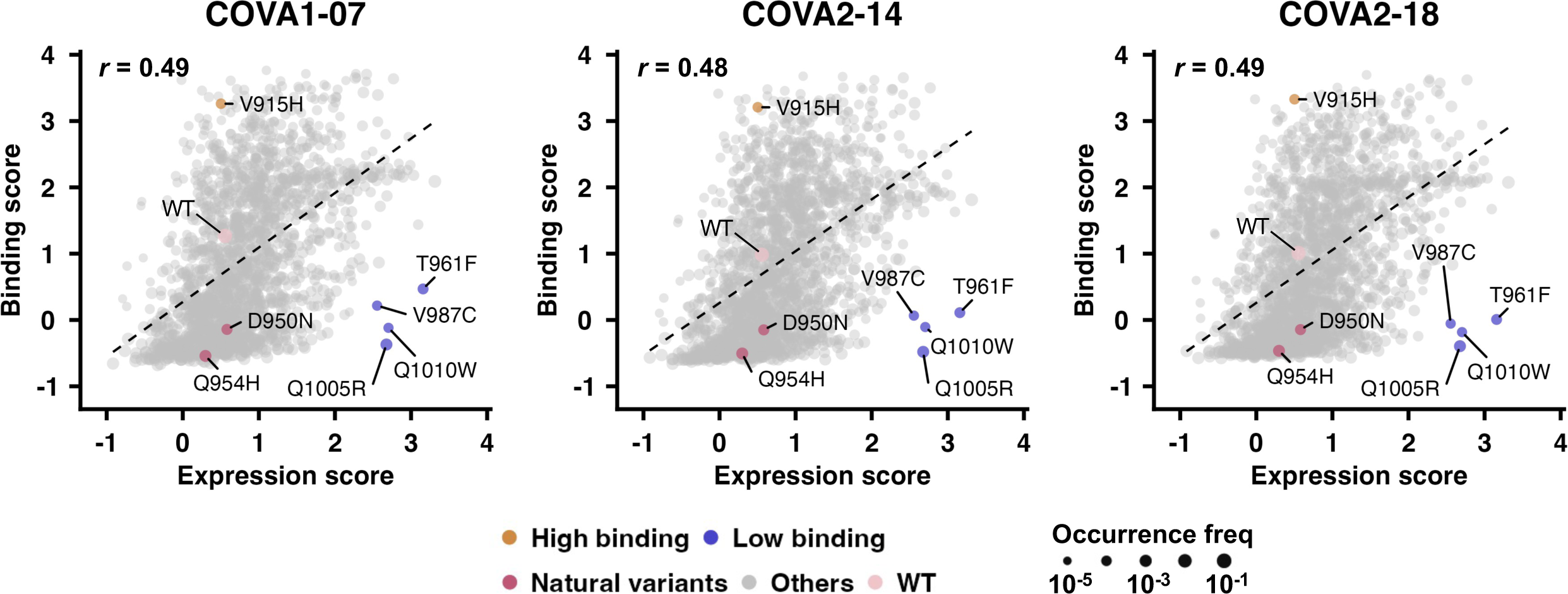
Identification and validation of public S2 HR1/CH antibody escape mutations. Plots of binding scores to **(A)** COVA1-07, **(B)** COVA2-14, and **(C)** COVA2-18 against expression scores. One datapoint represents one amino acid mutation in our mutant library. Expression scores were obtained from our previous study^24^.

To experimentally validate our findings, S protein bearing S2 mutations D950N, Q954H, T961F, V987C, Q1005R, and Q1010W were individually expressed by transient transfection of HEK293T cells, which were subsequently analyzed by flow cytometry using COVA1-07, COVA2-14, COVA2-18, and CC12.3 (**Figure S3**). Compared to wild type (WT), these six mutations have lower cell-surface binding activity to COVA1-07, COVA2-14, and COVA2-18, albeit to different extents (**Figure 3A-C**). T961F, Q1005R, and Q1010W weakened the cell-surface binding activity of these three antibodies by >80%, whereas D950N, Q954H, and V987C only weakened their binding activity by ∼40%, ∼25%, and ∼50%, respectively. In contrast, their cell-surface binding activity to the RBD antibody CC12.3 was similar to or higher than WT (**Figure 3D**). In this validation experiment, we also included mutation V915H as a control, which had a binding score of >3 against COVA1-07, COVA2-14, and COVA2-18 in our deep mutational scanning experiments, but a WT-like expression score (**Figure 2 and Figure 3A-D**). These results indicate that D950N, Q954H, T961F, V987C, Q1005R, and Q1010W can weaken the binding activity of IGHV1-69/IGKV3-11 S2 antibodies. Moreover, since D950N and Q954H can be observed in the natural variants of SARS-CoV-2^1^, our results also suggest that antigenic drift occurs in the S2 domain.

**Figure 3.**
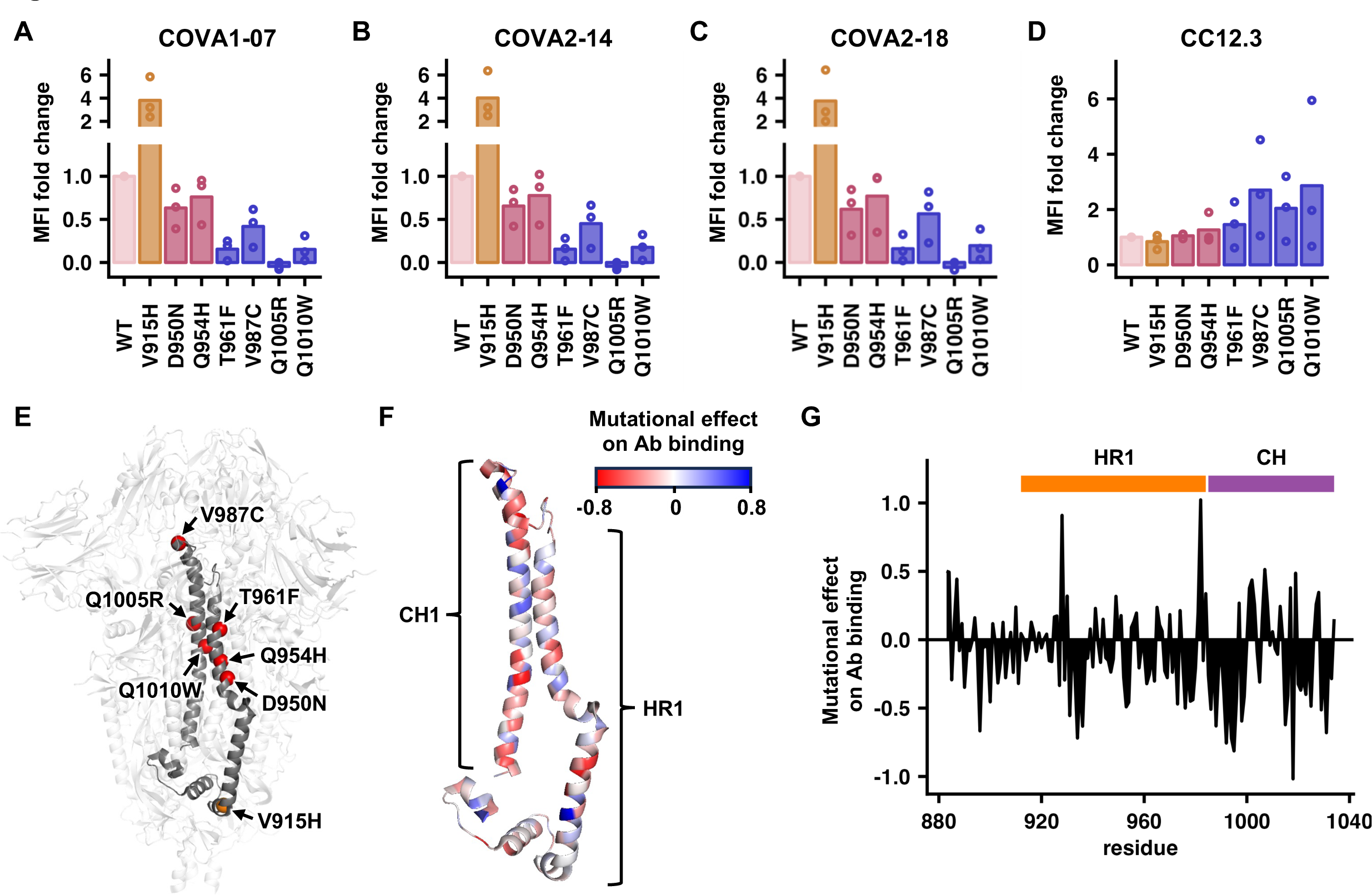
Experimental validation of mutations that influence binding to IGHV1-69/IGKV3-11 S2 antibodies (A-D) The binding of individual mutants to **(A)** COVA1-07 **(B)** COVA2-14, **(C)** COVA2-18, and **(D)** CC12.3 was individually analyzed using flow cytometry. For each mutant, fold change of median fluorescence intensity (MFI) compared to wild type (WT) is shown (**see Methods**). **(E)** The locations of D950N, Q954H, T961F, V987C, Q1005R, and Q1010W are highlighted as red spheres on one protomer of the SARS-CoV-2 S structure (PDB 6VXX)^43^. The location of V915H is highlighted as an orange sphere. Our mutant library contained mutations from residues 883 to 1034, which are in gray on one protomer. Three independent biological replicates were performed. **(F)** Mutational effect on antibody binding is shown on the SARS-CoV-2 S structure (PDB 6VXX)^43^. Only residues 883 to 1034 of one protomer is shown. **(G)** Mutational effect on antibody binding is plotted against the residue position.

### Spatially diverse S2 residues modulate binding to IGHV1-69/IGKV3-11 antibodies

S2 mutations that were validated to influence the binding activity of IGHV1-69/IGKV3-11 antibodies were spatially distributed widely (**Figure 3E**). For example, V987C, which decreased the binding to IGHV1-69/IGKV3-11 antibodies, resided at the S2 apex. In contrast, V915H, which increased the binding to IGHV1-69/IGKV3-11 antibodies, was at the bottom of the S2 domain. Other experimentally validated mutations, namely D950N, Q954H, T961F, Q1005R, and Q1010W, were located at the center of the S2 domain.

We further performed a systematic analysis using our deep mutational scanning data. To correct for the effect of S expression level on cell-surface binding activity, we computed an adjusted binding score, which represented the residual of a linear regression model of binding score on expression score. Subsequently, for a given S2 residue, the mutational effect on antibody binding was computed as the average adjusted binding score for all mutations at that residue (**see Methods**). A positive mutational effect on antibody binding indicated that mutations tended to increase binding to IGHV1-69/IGKV3-11 antibodies, whereas negative indicated that mutations tended to decrease binding. Consistent with diverse spatial distribution of the validated mutations, residues with strong mutational effects on antibody binding, either positive or negative, were spread across the S2 domain (**Figure 3F-G**). Similarly, a previous study has shown that the binding activity of COV2-2164 and CnC2t1p1 B10, which belong to the same IGHV1-69/IGKV3-11 clonotype as COVA1-07, COVA2-14, and COVA2-18, can be abolished by spatially distinct mutations in the S2 (K814A, I980A, R995A, and Q1002A)^19^. Therefore, the binding activity of IGHV1-69/IGKV3-11 antibodies can be modulated by S2 mutations that are outside of the epitope.

Since the epitope of these IGHV1-69/IGKV3-11 antibodies is exposed only when the S protein is in open conformations^18^, any mutations that alter the conformational dynamics of the S protein may affect the binding to IGHV1-69/IGKV3-11 antibodies. For example, HexaPro, which consists of six proline mutations to stabilize the prefusion conformation, was previously shown to dramatically decrease the on-rate of the IGHV1-69/IGKV3-11 antibodies^18^. Consistently, S2 mutations T961F and Q1005R, which were validated to weaken the binding to IGHV1-69/IGKV3-11 antibodies (**Figure 3A-D**), were previously shown to be fusion-incompetent^24^. Previous studies also hinted that the natural mutations D950N and Q954H altered the conformational dynamics of the S protein^27,28^ – D950N was shown to slightly promote membrane fusion^27^, whereas Q954H was shown to favor a kinked conformation of HR1^28^. These observations could explain the diverse spatial distribution of S2 residues that influence binding to IGHV1-69/IGKV3-11 antibodies.

### S2 mutation D950N weaken the *in vivo* protection activity of COVA2-18

At the end, we aimed to test if the *in vivo* protection activity of IGHV1-69/IGKV3-11 antibodies can be escaped by natural mutations in the S2 domain. Consequently, we focused on D950N, which was a natural mutation that weakened binding to IGHV1-69/IGKV3-11 antibodies (**Figure 3A-D**). Mutation D950N was introduced into recombinant SARS2-N501Y_MA30_ (rSARS2-N501Y_MA30_). Mice infected with rSARS2-N501Y_MA30_ WT were partially protected by COVA2-18 that trended towards statistical significance (*P* = 0.11, **Figure 4A**). In contrast, COVA2-18 was not protective in mice infected with rSARS2-N501Y_MA30_ D950N (*P* = 0.92, **Figure 4B**). These data imply that mutations in S2 may contribute to the natural escape of protective antibody response.

**Figure 4.**
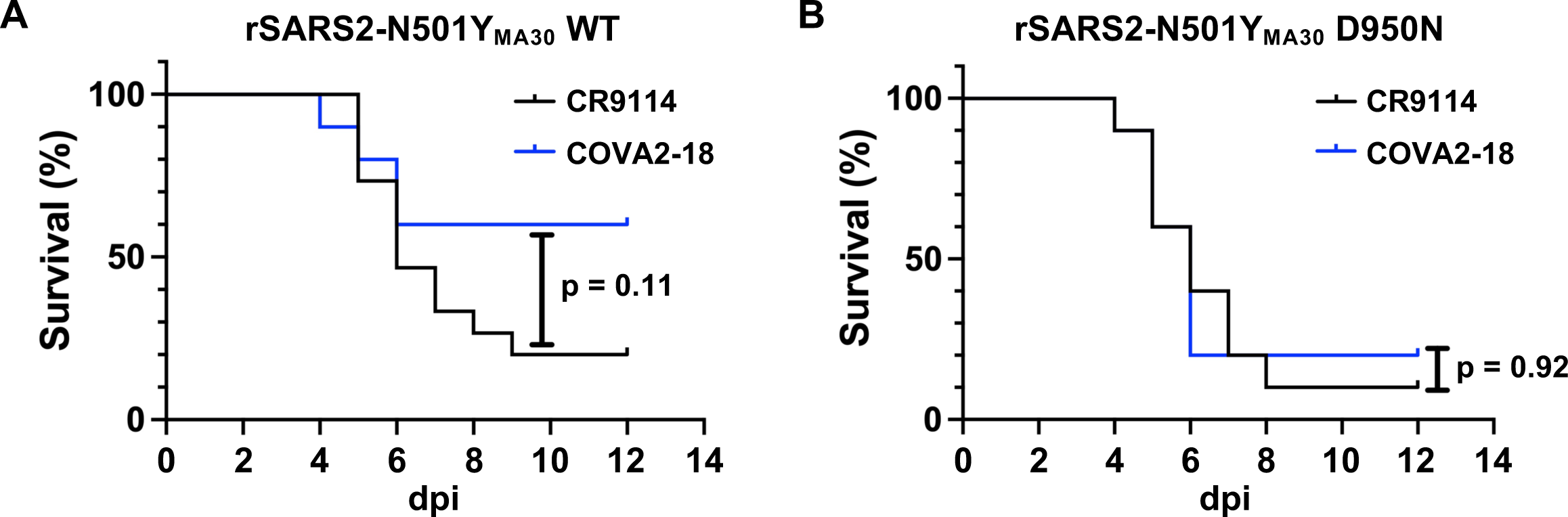
S2 mutation D950N escaped the in vivo protection activity of COVA2-18. BALB/c mice were intravenously injected with 200 μg of CR9114 (black) or COVA2-18 (blue) 16 hours before infection. Mice treated with antibodies were subsequently infected intranasally with 1000 PFU of **(A)** rSARS2-N501Y_MA30_ WT or **(B)** rSARS2-N501Y_MA30_ encoding the S2 mutation D950N mutation. Kaplan-Meier survival curves are shown. *n* = 10 for all groups, except for the group treated with CR9114 in **(A)**, where *n* = 15.

## DISCUSSION

Public antibodies against viral antigens such as influenza hemagglutinin, HIV envelope protein, and SARS-CoV-2 S have been documented^29–31^. Since these antibodies are found in many individuals, public antibodies can exert a positive selection pressure on virus at the population level, which in turn promote antigenic drift. For example, natural mutations K417N/T in the S of many SARS-CoV-2 variants abrogate binding to public RBD antibodies encoded by IGHV3-53/3-66^32–34^. Our study here suggests a similar phenomenon in the relatively conserved core fusion machinery of SARS-CoV-2 S protein. Specifically, we show that natural S2 mutations D950N and Q954H can weaken the binding to public antibodies encoded by IGHV1-69/IGKV3-11. Our results further suggest that the *in vivo* protection activity of these IGHV1-69/IGKV3-11 antibodies can be affected by natural S2 mutations. Therefore, while neutralizing antibodies are known to exert positive selection pressure on SARS-CoV-2^1,4,5^, non-neutralizing antibodies that are protective, such as IGHV1-69/IGKV3-11 S2 antibodies^18^, may do the same but to a lesser extent.

A previous study has shown that the binding of IGHV1-69/IGKV3-11 antibodies to the S2 apex requires an open conformation of the S protein^18^. These observations imply that the epitope of IGHV1-69/IGKV3-11 S2 antibodies is exposed for binding when the S protein is transitioning from prefusion to postfusion conformations. If the epitope is only available during an intermediate state between prefusion and postfusion conformations of the S protein, any mutations that decrease the half-life of such intermediate state will weaken the binding to IGHV1-69/IGKV3-11 S2 antibodies. These mutations can either stabilize the prefusion conformation to prevent transition to the intermediate state or accelerate the transition from the intermediate state towards the postfusion conformation. Consistently, mutations that weaken the binding to IGHV1-69/IGKV3-11 S2 antibodies include those that abolish fusion activity (e.g. T961F and Q1005R)^24^ and enhance fusion activity (e.g. D950N)^27^. Nevertheless, the conformation dynamics of S protein during the fusion process remains largely elusive. In addition, a high-resolution structure of an IGHV1-69/IGKV3-11 S2 antibody in complex with S2 is lacking. These limitations prevent a detailed mechanistic understanding of mutations that weaken the binding to IGHV1-69/IGKV3-11 S2 antibodies.

As SARS-CoV-2 continues to evolve in the human population and other coronavirus strains remain a pandemic threat, there is an urge to develop a more universal coronavirus vaccine^35^. Its feasibility is substantiated by the discovery of broadly protective antibodies to the highly conserved S2 domain^9–17^. However, our results suggest that escape mutations against S2 antibodies can emerge in naturally circulating SARS-CoV-2 variants. A previous study has shown that S2 mutation D796H, which locates near the base of the S protein and emerged in a chronically infected patient, reduce sensitivity to neutralization by convalescent plasma^36^. More recently, a deep mutational scanning study of the full SARS-CoV-2 S protein has also identified natural mutations that can escape antibodies to S2 stem helix^37^. Consequently, potential escape mutations may impose a challenge for S2-based vaccine development^38–40^.

## METHODS

### Cell culture

Human embryonic kidney 293T (HEK293T) cells (ATCC) were grown and maintained at 37 °C, 5% CO_2_ and 95% humidity in D10 medium: Dulbecco’s modified Eagle medium (DMEM) with high glucose (Gibco) supplemented with 10% v/v fetal bovine serum (FBS; Gibco), 1× non-essential amino acids (Gibco), 1× GlutaMAX (Gibco), 100 U/mL penicillin and 100 μg/mL streptomycin (Gibco). HEK293T landing pad cells were grown and maintained in D10 medium supplemented with 2 μg/mL doxycycline (Thermo Scientific) at 37 °C, 5% CO_2_ and 95% humidity. Expi293F suspension cells (Gibco) were grown and maintained in Expi293 expression medium (Gibco) at 37 °C, 125 rpm, 8% CO_2_ and 95% humidity according to the manufacturer’s instructions.

### Plasmid construction

The S2 HR1/CH mutant library was constructed in our previous study. For experimental validation, codon-optimized oligonucleotide encoding S (GenBank: MN908947.3) with the PRRA motif in the furin cleavage site deleted was cloned into a phCMV3 vector. Site-directed mutagenesis was performed using PrimeSTAR Max DNA polymerase (Takara Bio) with the following settings: 98 °C for 10 s, 22 cycles of (98 °C for 10 s, 55 °C for 5 s, 72 °C for 45 s), 72 °C for 45 s. The PCR product was digested with DpnI (NEB) for 2 h at 37 °C and transformed into chemically competent DH5α *Escherichia coli*.

### Antibody expression and purification

The heavy and light chain sequences of COVA1-07, COVA2-14, COVA2-18, CC12.3, and CR9114 were synthesized commercially (Integrated DNA Technologies), amplified via PCR, and cloned into a phCMV3 vector in an IgG1 format. Plasmids encoding heavy and light chains were transfected into Expi293F cells in a 2:1 mass ratio using an ExpiFectamine 293 transfection kit (Gibco). Six days post-transfection, the supernatant was harvested via centrifugation of cell suspension at 4 °C and 4500 × *g* for 30 min. The supernatant was clarified using a polyethersulfone membrane filter with a 0.22 μm pore size (Millipore).

CaptureSelect CH1-XL beads (Thermo Scientific) were washed thrice with MilliQ H_2_O and resuspended in 1× PBS. The clarified supernatant was incubated with washed beads at 4 °C overnight with gentle rocking. Flowthrough was subsequently collected, and beads washed once with 1× PBS. Beads were incubated in 60 mM sodium acetate, pH 3.7 for 10 min at 4 °C for elution of the antibodies. Antibodies were further purified by size-exclusion chromatography using a Superdex 200 XK 16/100 column in 1× PBS. Fractions corresponding to ∼150 kDa were pooled and concentrated using a centrifugal filter unit with a 30 kDa molecular weight cut-off (Millipore) via centrifugation at 4000 × *g* and 4 °C for 15 min. Antibodies were stored at 4 °C.

### Fluorescence-activated cell sorting

HEK293T landing pad cells expressing the S2 HR1/CH mutant library of S as constructed previously^24^ were harvested and centrifuged at 300 × *g* for 5 min at 4 °C. Supernatant was discarded. Cells were resuspended in ice-cold FACS buffer. Cells were incubated with 3 μg/mL of COVA1-07, 3 μg/mL of COVA2-14, or 10 μg/mL of COVA2-18 for 1 h at 4 °C with gentle rocking. Subsequently, cells were washed once, and resuspended with ice-cold FACS buffer. Cells were incubated with 2 μg/mL of PE anti-human IgG Fc. Cells were washed once, resuspended in ice-cold FACS buffer, and filtered through a 40 μm strainer. Cells were sorted via a three-way sort using a BigFoot spectral cell sorter (Invitrogen) according to levels of PE fluorescence at 4 °C. Cells with no PE fluorescence were sorted into “bin 0”. Cells with PE at similar levels as WT were sorted into “bin 1”. Cells with PE at higher levels than WT were sorted into “bin 2”. Gating strategy is shown in **Figure S1**. Cells collected per bin for each replicate and antibody are listed in **Table S1**.

### Post-sorting genomic DNA extraction

Cell pellets were obtained via centrifugation at 300 × *g* for 15 min at 4 °C, and the supernatant was discarded. Genomic DNA was extracted using a DNeasy Blood and Tissue Kit (Qiagen) following the manufacturer’s protocols with a modification, where resuspended cells were lysed at 56 °C for 30 min instead of 10 min.

### Next-generation sequencing

After genomic DNA extraction, the region of interest spanning the HR1 and CH was amplified via PCR using 5′-CAC TCT TTC CCT ACA CGA CGC TCT TCC GAT CTA CAT CTG CCC TGC TGG CCG GCA CA-3′ and 5′-GAC TGG AGT TCA GAC GTG TGC TCT TCC GAT CTG CAA AAG TCC ACT CTC TTG CTC TG-3′ as forward and reverse primers, respectively. A maximum of 500 ng of genomic DNA was used as template per reaction. In total, a maximum of 4 μg per bin per replicate was used as template (i.e. 8 reactions). PCR was performed using KOD DNA polymerase (Takara Bio) with the following settings: 95 °C for 2 min, 25 cycles of (95 °C for 20 s, 56 °C for 15 s, 68 °C for 20 s), 68 °C for 2 min, 12 °C hold. After PCR, all eight 50 μL reactions per bin per replicate were mixed. 100 μL of product per bin per replicate was used for purification using a PureLink PCR Purification kit (Invitrogen). Subsequently, 10 ng of the purified PCR product per bin per replicate was appended with Illumina barcodes via PCR using primers: 5′-AAT GAT ACG GCG ACC ACC GAG ATC TAC ACX XXX XXX XAC ACT CTT TCC CTA CAC GAC GCT-3′, and 5′-CAA GCA GAA GAC GGC ATA CGA GAT XXX XXX XXG TGA CTG GAG TTC AGA CGT GTG CT-3’. Positions annotated by an “X” represented the nucleotides for the index sequence. This PCR was performed using KOD DNA polymerase with the following settings: 95 °C for 2 min, 9 cycles of (95 °C for 25 s, 56 °C for 15 s, 68 °C for 20 s), 68 °C for 2 min, 12 °C hold. Barcoded products were mixed and sequenced with a MiSeq PE300 v3 flow cell (Illumina).

### Analysis of next-generation sequencing data

Forward and reverse reads were merged using PEAR^41^. Forward reads were translated and matched to the corresponding mutant. Counts for each bin for each replicate were tabulated. A pseudocount of 1 was added to all counts. For each replicate, the frequency of each mutant was calculated as the count of that mutant divided by the total number of counts in that bin:

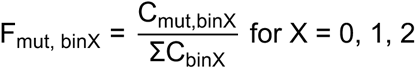

For each replicate, the binding score of each mutant (B_mut_) was calculated using:

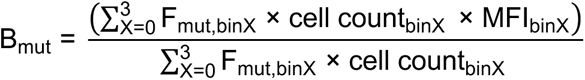

Cell counts for bin0, bin1, and bin2 had a ratio of approximately 100:10:1. As a result, cell count_bin0_, cell count_bin1_, and cell count_bin2_ were set as 100, 10, and 1, respectively. Similarly, the median fluorescence intensities (MFI) for bin0, bin1, and bin2 had a ratio of approximately 1:50:500. Therefore, MFI_bin0_, MFI_bin1_, and MFI_bin2_ were set as 1, 50, and 500, respectively. Subsequently, the binding score of all mutants was scaled to obtain the adjusted binding score 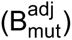 such that the means of adjusted binding scores of nonsense mutations and silent mutations equal 0 and 1, respectively, using the following equation:

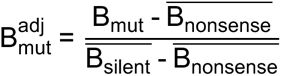

where 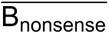 and 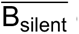 correspond to the means of non-adjusted binding scores of nonsense mutations and silent mutations, respectively.

### Flow cytometry

To validate and quantify surface expression of WT S and its mutants, flow cytometry was performed. 4 × 10^6^ HEK293T cells were seeded on wells of 6-well plates and incubated overnight at 37 °C. Then, cells were transfected with 2 μg of plasmid encoding WT S or the indicated mutant using Lipofectamine 2000 (Invitrogen) following the manufacturer’s protocol. At 20 hours post-transfection, cells were harvested and resuspended in ice-cold FACS buffer (2% v/v FBS, 50 mM EDTA in DMEM supplemented with high glucose, L-glutamine and HEPES, without phenol red [Gibco]). Cells were incubated with 5 μg/mL of RBD antibody CC12.3^25^ for 1 h at 4 °C with gentle rocking. Then, cells were washed once, and resuspended with ice-cold FACS buffer. Cells were incubated with 2 μg/mL of phycoerythrin (PE) anti-human IgG Fc (Clone M1310G05, BioLegend). Cells were washed once, resuspended in ice-cold FACS buffer, and filtered through a 40 μm strainer. Subsequently, cells were analyzed using an Accuri C6 flow cytometer (BD Biosciences). Gating strategy is shown in **Figure S3**.

Binding of WT or mutant S to the S2 HR1 public antibodies was validated and quantified via flow cytometry. The same protocol was followed as above except that cells were incubated with 3 μg/mL of COVA1-07, 3 μg/mL of COVA2-14, or 10 μg/mL of COVA2-18. The concentrations of these antibodies were determined via titration using HEK293T landing pad cells stably expressing WT S on their surface. Median fluorescence intensity (MFI) values were calculated by plotting data in FCS Express 6 software (De Novo Software). Fold change in MFI was calculated using the following equation:

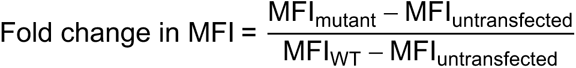

### Introduction of S mutations D950N and Q954H into rSARS2-N501Y_MA30_ BAC

rSARS2-N501Y_MA30_ BACs with D950N and Q954H mutations in S were individually generated using lambda red recombination system with I-SceI homing endonuclease as described previously^42^. In brief, forward and reverse primers carrying overlapping SARS-CoV-2 sequences with D950N or Q954H mutation followed by a sequence complementary to the target plasmid (pEP-KanS) were designed. PCR fragments with overlapping ends carrying SARS-CoV-2 sequence flanking kanamycin resistance marker was amplified with the primers described using pEP-KanS as template. PCR fragments were purified using a PureLink Quick Gel Extraction and PCR Purification Combo Kit (Invitrogen). Purified PCR fragments were electroporated into the GS1783 strain of E. coli carrying the rSARS2-N501Y_MA30_ BAC. The kanamycin markers were removed by arabinose induction of I-SceI cleavage followed by homologous recombination of the overlapping ends. Successful recombinants were selected by replica plating for the loss of kanamycin resistance. Recombinants were purified and sequenced to confirm the introduction of the mutation into the BAC.

The primer sequences used for the D950N mutation are as follows:

Forward: 5’-TCC ACA GCA AGT GCA CTT GGA AAA CTT CAA aac GTG GTC AAC CAA AAT GCA CAA GCT TTA AAC ACG CTT GTT AAA AGG ATG ACG ACG ATA AGT AG-3’
Reverse: 5’-ATC ATT TAA AAC ACT TGA AAT TGC ACC AAA ATT GGA GCT AAG TTG TTT AAC AAG CGT GTT TAA AGC TTG TGC ATT CAA CCA ATT AAC CAA TTC TGA TTA G-3’

The primer sequences used for the Q954H mutation are as follows:

Forward: 5’-ACT TGG AAA ACT TCA AGA TGT GGT CAA Cca cAA TGC ACA AGC TTT AAA CAC GCT TGT TAA ACA ACT TAG CTC CAA AGG ATG ACG ACG ATA AGT AG-3’
Reverse: 5’-AGA CGT GAA AGG ATA TCA TTT AAA ACA CTT GAA ATT GCA CCA AAA TTG GAG CTA AGT TGT TTA ACA AGC GTG TTT CAA CCA ATT AAC CAA TTC TGA TTA G-3’

Mutations are in lower-case letters. Sequences complementary to pEP-KanS are underlined. BAC sequences were validated by whole BAC sequencing.

### Rescue of BAC-derived virus and virus propagation

2 μg of rSARS2-N501Y_MA30_ BAC were separately transfected into Vero cells (ATCC) with Lipofectamine 3000 (Invitrogen) in a 6-well plate according to manufacturer’s protocol. Cells were monitored daily for cytopathic effects (CPE). Viruses in the supernatant were harvested when CPE was >50% and stored at -80 °C until used. Viruses were further passaged in Vero cells expressing TMPRSS2 in DMEM supplemented with 10% FBS. Virus sequences were confirmed by next-generation sequencing. Virus titers were determined by plaque assay. Briefly, viruses were serially diluted in DMEM. Twelve-well plates of Vero cells were inoculated at 37 °C in 5% CO_2_ for 1 h and gently rocked every 15 min. After removing the inocula, plates were overlaid with 0.6% agarose containing 2% FBS. After 3 days, overlays were removed, and plaques visualized by staining with 0.1% crystal violet. Viral titers were quantified as PFU/mL.

### *In vivo* virus challenge

8-10 week old female BALB/c mice were used in this study. Mice were anesthetized with ketamine-xylazine and infected intranasally with 5,000 PFU of the non-recombinant virus or 1,000 PFU of the recombinant virus in 50 μl of Dulbecco’s Modified Eagle Medium (DMEM) (Gibco). For antibody treatment, mice were injected with 200 μg of antibody (CR9114, COVA1-07, COVA2-14, COVA2-18) diluted in 200 μL of PBS through intravenous route one day before infection. All work with SARS-CoV-2 was performed in the University of Iowa’s Biosafety level (BSL3) laboratories. All animal studies were approved by the University of Iowa Animal Care and Use Committee and meet stipulations of the Guide for the Care and Use of Laboratory Animals.

### Statistics

All indicated statistical tests were performed using R.

## DATA AVAILABILITY

Raw reads from deep sequencing data can be accessed at BioProject accession PRJNA888135.

## CODE AVAILABILITY

Custom code to analyze deep sequencing data for escape mutations is available at https://github.com/nicwulab/SARS2_HR1_DMS_Abs.

## AUTHOR CONTRIBUTIONS

T.J.C.T. and N.C.W. conceived and designed the study. T.J.C.T. performed flow cytometry. T.J.C.T. and N.C.W. analyzed deep sequencing data. T.J.C.T. and R.L. performed antibody expression and purification. A.O. and L.R.W. performed the *in vivo* experiments. K.A.M. provided the HEK293T landing pad cell line. S.P. and N.C.W. provided resources and support. T.J.C.T., L.R.W., and N.C.W. wrote the paper and all authors reviewed and edited the paper.

## ETHICAL DECLARATIONS

N.C.W. serves as a consultant for HeliXon. All authors declare no other competing interests.

## ACKNOWLEDGEMENTS

We thank the Roy J. Carver Biotechnology Center at the University of Illinois Urbana-Champaign for assistance with fluorescence-activated cell sorting and performing deep sequencing, and Meng Yuan for helpful discussion. This work was supported by National Institutes of Health K99 AI170996 (L.R.W.), R01 AI165475 (N.C.W.) and the Searle Scholars Program (N.C.W.).

## Notes

https://www.ncbi.nlm.nih.gov/bioproject/?term=PRJNA888135

https://github.com/nicwulab/SARS2_HR1_DMS_Abs

