## Supplementary Information for "Evidence of antigenic drift in the fusion machinery core of SARS-CoV-2 spike"

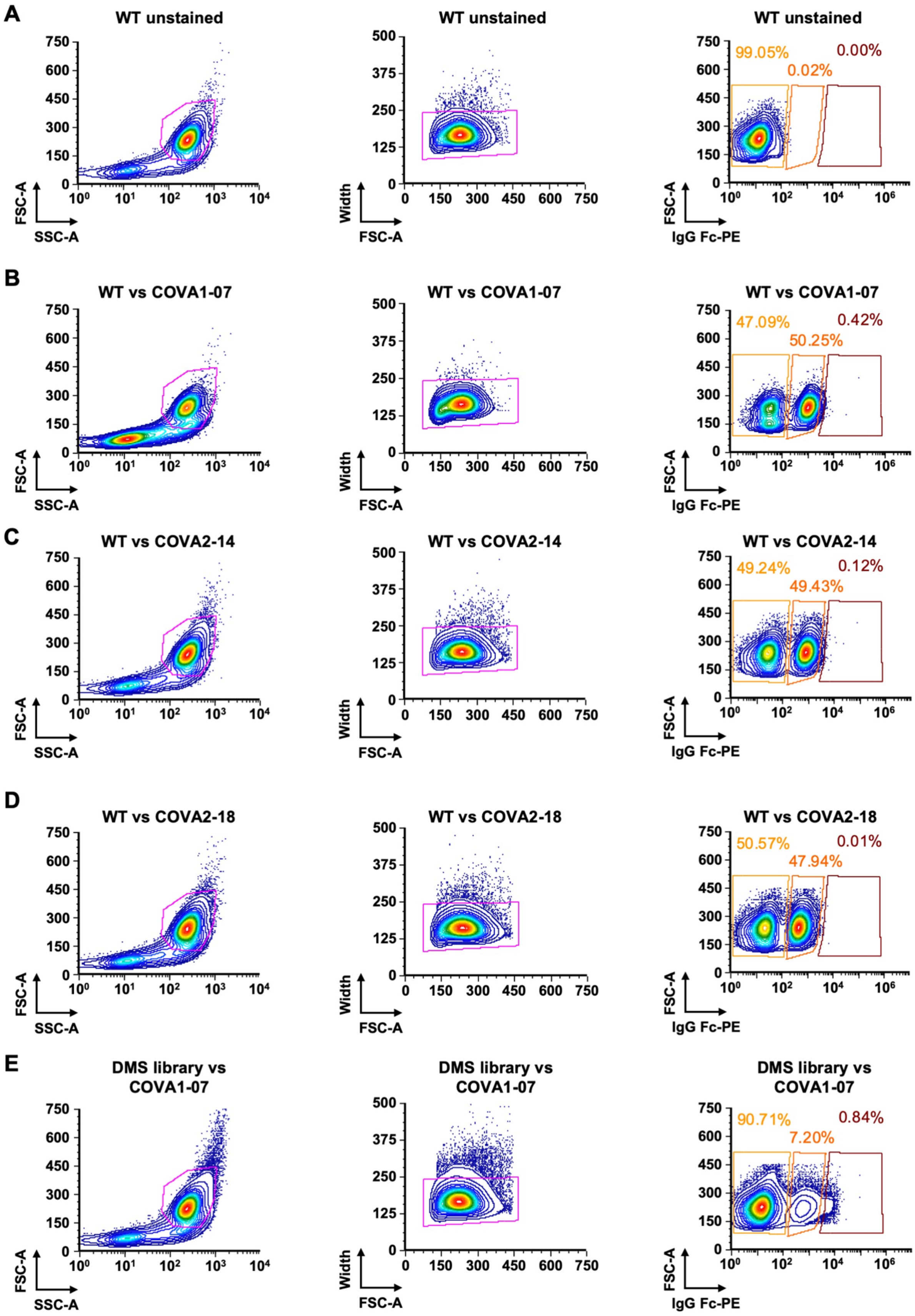
A
WT unstained
FSC-A
SSC-A
Width
FSC-A
99.05%
0.02%
0.00%
IgG Fc-PE
B
WT vs COVA1-07
47.09%
50.25%
0.42%
C
WT vs COVA2-14
49.24%
49.43%
0.12%
D
WT vs COVA2-18
50.57%
47.94%
0.01%
E
DMS library vs COVA1-07
90.71%
7.20%
0.84%

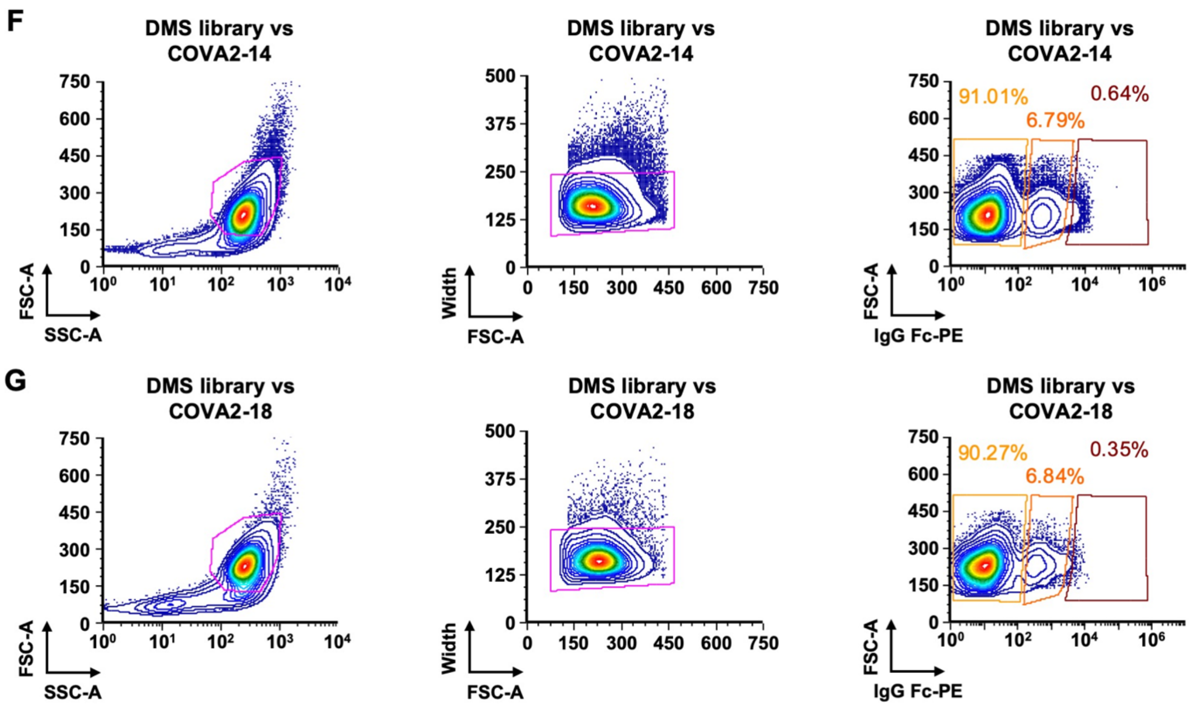


**Figure S1. Gating strategy for FACS of DMS library. (A)** Flow cytometry plots of the parental HEK293T landing pad cells without expression of SARS-CoV-2 spike. Primary antibody used was COVA1-07. **(B-D)** Flow cytometry plots of HEK293T landing pad cells stably expressing WT SARS-CoV-2 spike. Primary antibodies used were **(B)** COVA1-07, **(C)** COVA2-14, and **(D)** COVA2-18. **(E-G)** Flow cytometry plots of HEK293T landing pad cells stably expressing the DMS library. Primary antibodies used were **(E)** COVA1-07, **(F)** COVA2-14, and **(G)** COVA2-18.

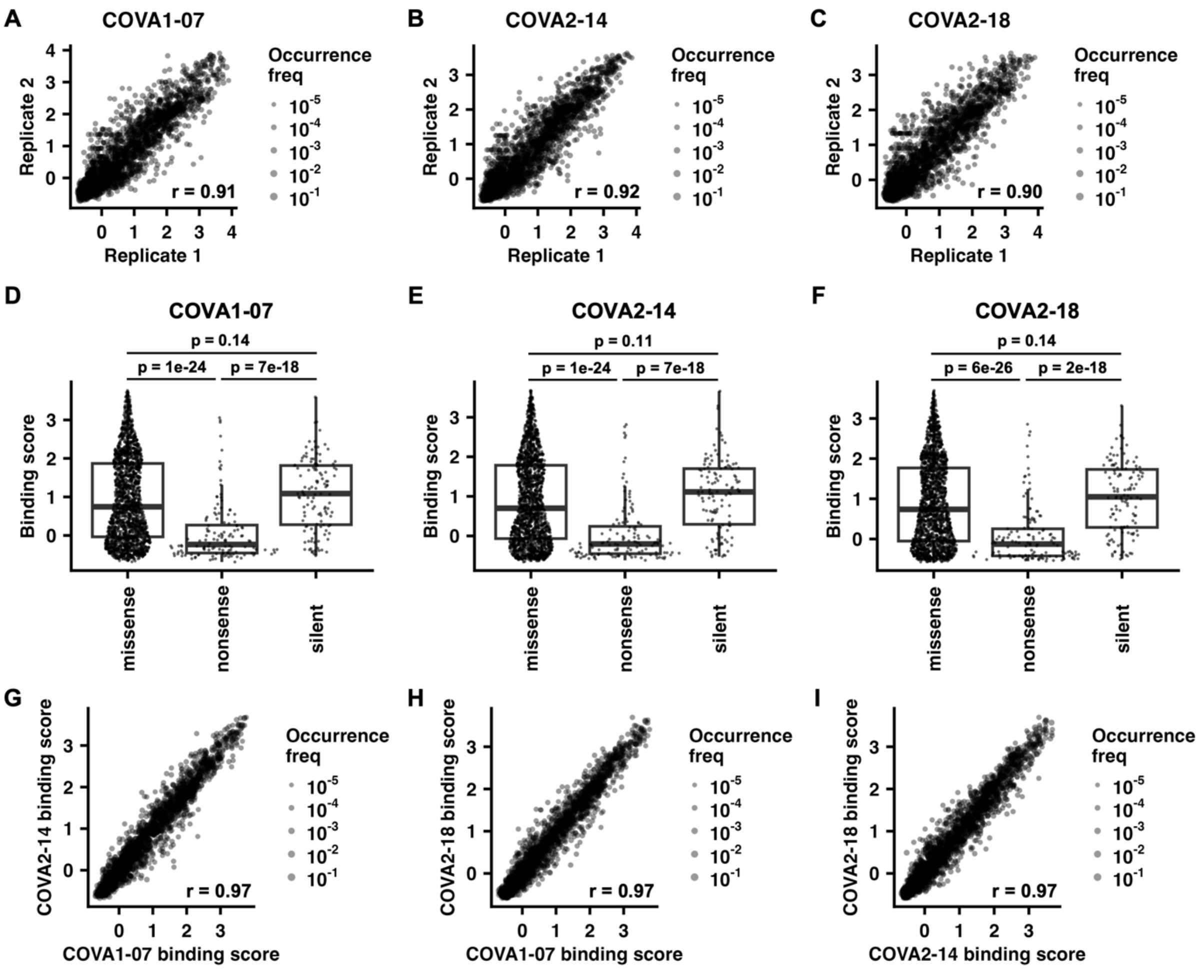


**Figure S2. Reproducibility and quality of the deep mutational scanning data. (A-C)** Correlation of binding scores to **(A)** COVA1-07, **(B)** COVA2-14, and **(C)** COVA2-18 between two biological replicates. Pearson's correlation coefficient, *r*, for each plot is indicated. **(D-F)** Box plots showing distribution of average binding scores to **(D)** COVA1-07, **(E)** COVA2-14, and **(F)** COVA2-18 according to different mutation classes (missense, nonsense, and silent). Two-tailed Student's *t* tests were performed, and *p*-values are shown. **(G-I)** Correlation plots between the binding scores to **(G)** COVA1-07 and COVA2-14, **(H)** COVA1-07 and COVA2-18, and **(I)** COVA2-14 and COVA2-18.

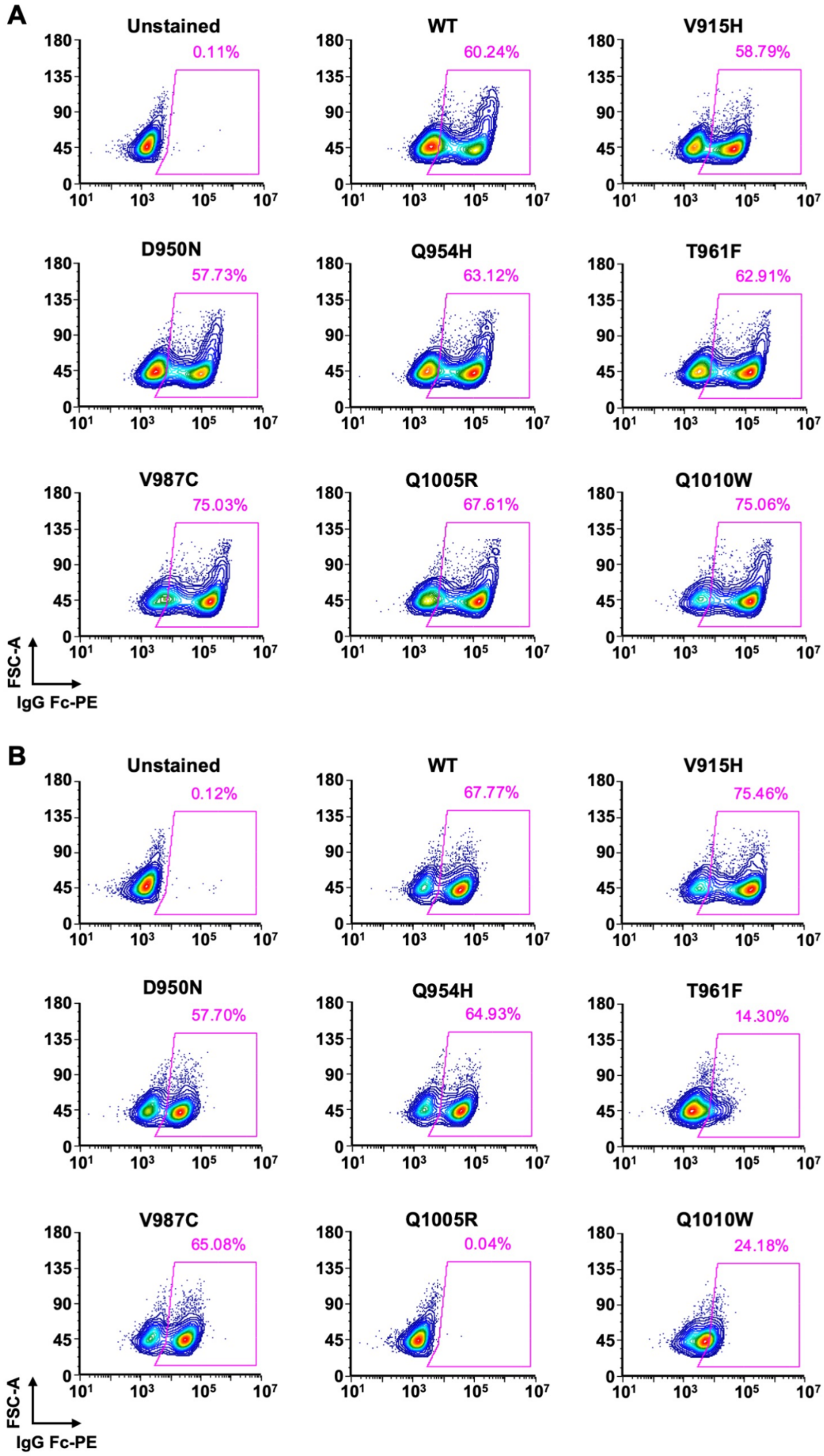

A
Unstained
0.11%
WT
60.24%
V915H
58.79%
D950N
57.73%
Q954H
63.12%
T961F
62.91%
V987C
75.03%
Q1005R
67.61%
Q1010W
75.06%
FSC-A
IgG Fc-PE
B
Unstained
0.12%
WT
67.77%
V915H
75.46%
D950N
57.70%
Q954H
64.93%
T961F
14.30%
V987C
65.08%
Q1005R
0.04%
Q1010W
24.18%
FSC-A
IgG Fc-PE

C

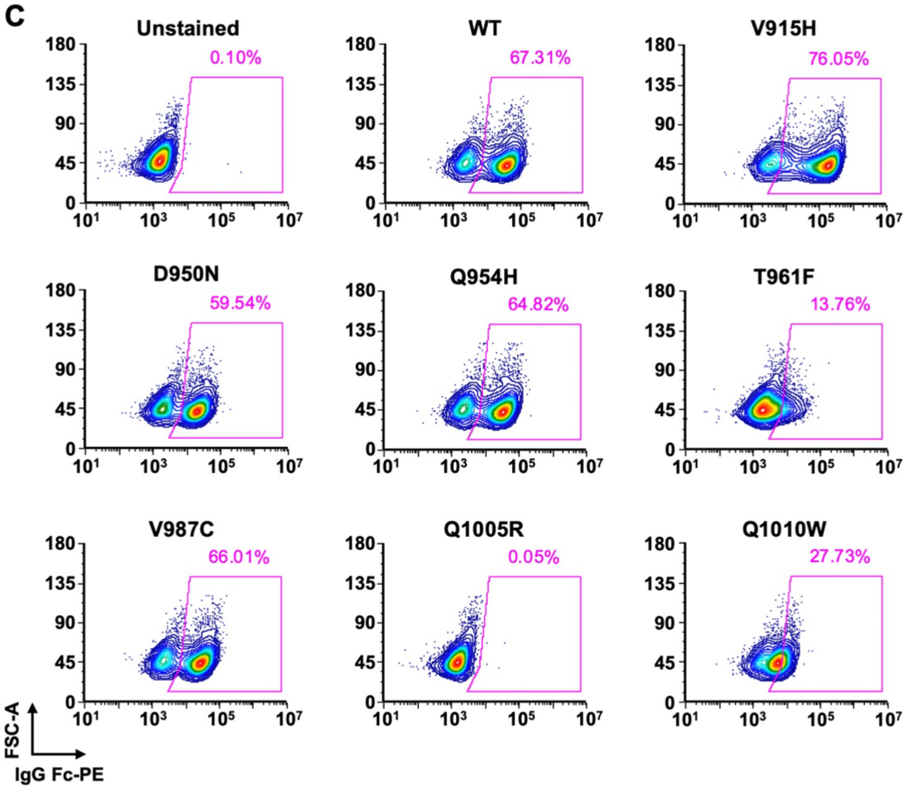

Unstained
0.10%
WT
67.31%
V915H
76.05%
D950N
59.54%
Q954H
64.82%
T961F
13.76%
V987C
66.01%
Q1005R
0.05%
Q1010W
27.73%
180
135
90
45
0
10^1
10^3
10^5
10^7
FSC-A
IgG Fc-PE


D

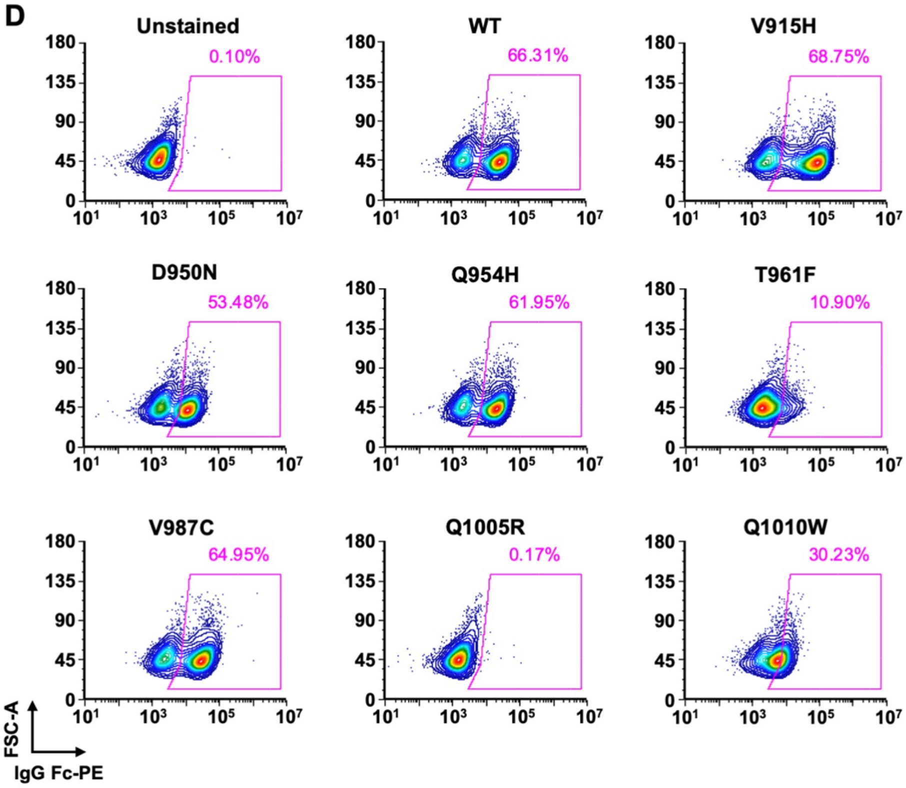

Unstained
0.10%
WT
66.31%
V915H
68.75%
D950N
53.48%
Q954H
61.95%
T961F
10.90%
V987C
64.95%
Q1005R
0.17%
Q1010W
30.23%
180
135
90
45
0
10^1
10^3
10^5
10^7
FSC-A
IgG Fc-PE

**Figure S3. Flow cytometry plots for validation of expression and binding. (A)** Cytometry plots for validation and quantification of surface expression of WT or mutant S using CC12.3, which is an RBD antibody [S1]. **(B-D)** Cytometry plots for validation and quantification of binding of WT or mutant S to **(B)** COVA1-07, **(C)** COVA2-14, or **(D)** COVA2-18.

**Table S1.** Number of cells sorted into each bin in fluorescence-activated cell sorting.

| COVA1-07 | | | |
|---|---|---|---|
| | **Bin 0** | **Bin 1** | **Bin 2** |
| **Replicate 1** | $3.69 \times 10^6$ | $5.65 \times 10^5$ | $1.11 \times 10^5$ |
| **Replicate 2** | $4.14 \times 10^6$ | $6.72 \times 10^5$ | $8.03 \times 10^4$ |
| **COVA2-14** | | | |
| | **Bin 0** | **Bin 1** | **Bin 2** |
| **Replicate 1** | $3.51 \times 10^6$ | $9.43 \times 10^5$ | $1.52 \times 10^5$ |
| **Replicate 2** | $3.98 \times 10^6$ | $5.64 \times 10^5$ | $9.10 \times 10^4$ |
| **COVA2-18** | | | |
| | **Bin 0** | **Bin 1** | **Bin 2** |
| **Replicate 1** | $3.22 \times 10^6$ | $5.90 \times 10^5$ | $1.38 \times 10^5$ |
| **Replicate 2** | $4.11 \times 10^6$ | $9.76 \times 10^5$ | $1.19 \times 10^5$ |
